# Expression level of the reprogramming factor NeuroD1 is critical for neuronal conversion efficiency from different cell types

**DOI:** 10.1101/2021.10.19.465051

**Authors:** Kanae Matsuda-Ito, Taito Matsuda, Kinichi Nakashima

## Abstract

Several transcription factors, including NeuroD1, have been shown to act as neuronal reprogramming factors (RFs) that induce neuronal conversion from somatic cells. However, it remains unexplored whether expression levels of RFs in the original cells affect reprogramming efficiency. Here, we show that the neuronal reprogramming efficiency from two distinct glial cell types, microglia and astrocytes, is substantially dependent on the expression level of NeuroD1: low expression failed to induce neuronal reprogramming, whereas elevated NeuroD1 expression dramatically improved reprogramming efficiency in both cell types. Moreover, even under conditions where NeuroD1 expression was too low to induce effective conversion by itself, combined expression of three RFs (Ascl1, Brn2, and NeuroD1) facilitated the breaking down of cellular barriers, inducing neuronal reprogramming. Thus, our results suggest that a sufficiently high expression level of RFs or alternatively their combinatorial expression, is the key to achieving efficient neuronal reprogramming from different cells.

**Highlights:**

▪ High expression of NeuroD1 is required for neuronal conversion.
▪ **Multiple infections with NeuroD1-expressing virus enhance neuronal reprogramming**
▪ **Combinatorial expression of NeuroD1 with other RFs facilitates neuronal conversion**

**eTOC blurb:** In this article, Matsuda-Ito *et al*. demonstrate that the efficacy of conversion into neurons from two distinct glial cells, microglia and astrocytes, depends on the NeuroD1 expression level. They also show that increased NeuroD1 expression alone enables efficient neuronal reprogramming in non-reactive astrocytes that were previously shown to be difficult to convert into neurons.

## Introduction

Lineage-specific transcription factors induce direct reprogramming of somatic cells to other cell types, such as neurons, without passing through a pluripotent stem cell state. In 2010, mouse fibroblasts were shown to be directly converted to neurons *in vitro* by forced expression of three transcription factors, Ascl1, Brn2, and Myt1l (Vierbuchen et al., 2010). More recently, by examining combinations of factors used for reprogramming, it has become possible to convert fibroblasts to neurons of specific subtypes, such as sensory and motor neurons (Masserdotti et al., 2016; Matsuda and Nakashima, 2021). Several groups have reported *in vivo* neuronal reprogramming into neurons from glial cells, including astrocytes, which become reactive after brain damage and eventually contribute to glial scar formation. For example, ectopic expression of Sox2, Neurog2, or NeuroD1 converts endogenous astrocytes to neurons in the mouse brain (Guo et al., 2014; Mattugini et al., 2019; Niu et al., 2013; Wu et al., 2020). Microglia, a type of glial cell, are the resident immune cells in the brain and are derived from primitive macrophages (Ginhoux et al., 2010). Microglia accumulate at the injured site to remove dead cells after brain injury, such as stroke, and consequently become the predominant cell type in the ischemic core region (Annunziato et al., 2013). We have previously shown that microglia can be converted into neurons both *in vitro* and *in vivo* by the ectopic expression of lentivirus-encoded NeuroD1 (Matsuda et al., 2019). Although *in vivo* neuronal reprogramming from these two glial cells holds great promise as a therapeutic strategy, further improvement of neuronal reprogramming efficiency is warranted to supply sufficient numbers of new neurons for complete functional recovery from neurological injury and diseases.

In a previous report, single-cell RNA sequencing analysis indicated that high but not low expression of Ascl1 induced the expression of neuronal marker genes in fibroblasts (Treutlein et al., 2016), implying a correlation between transgene expression level and the attainment of neuronal reprogramming. However, it has not been extensively studied how the conditions in which reprogramming factors (RFs) are expressed influence neuronal reprogramming efficiency. Here, we examined neuronal reprogramming from microglia and astrocytes under conditions of different expression levels of NeuroD1 in these two glial cell types. In contrast to the higher expression, when we decreased the expression of NeuroD1 by reducing doxycycline (Dox) concentration, the neuronal conversion from microglia was dramatically diminished. On the other hand, increasing the NeuroD1 expression level by repeated lentiviral infections (2 times) improved neuronal reprogramming efficiency from microglia. Moreover, multiple NeuroD1-expressing viral infections (3 times) enabled neuronal reprogramming from non-reactive (NR-) astrocytes that were previously shown to be difficult to convert into neurons with a single infection (Matsuda et al., 2019). We also found that the combined expression of three RFs, Ascl1 and Brn2 together with NeuroD1, efficiently induced neuronal reprogramming, even when their expression was low. Taking these observations together, we believe that our study offers efficient strategies to reprogram neurons from glial cells and will contribute to accelerating the development of therapeutic applications for brain injury and diseases.

## Results

### NeuroD1 expression level-dependent changes in reprogramming efficiency

To investigate whether the efficiency of microglia-to-neuron (MtN) reprogramming is influenced by the expression level of NeuroD1, we used a lentivirus expressing NeuroD1 under the control of the Dox-inducible tetracycline response element promoter. We isolated CD68-, Iba1-, and Tmem119-positive (CD68^+^ Iba1^+^ Tmem119^+^) microglia from the 1-day-old mouse cortex with the same high purity as in our previous study (Matsuda et al., 2019) (Figure 1A) and added an equal amount of virus to each microglial culture dish, but treated them with different doses of Dox. At 7 days post treatment (dpt), we observed the Dox-dose dependent appearance of EGFP^+^ cells (Figures 1B and 1C). We next examined MtN conversion efficiency in the individual dishes based on the proportion of βIII-tubulin^+^/EGFP^+^ cells among Hoechst^+^ total cells and found that the efficiency was greatly decreased by reducing the Dox concentration (Figure 1D). Consistent with this, although no further increase of *NeuroD1* expression was observed at 2 μg/mL of Dox compared to 1 μg/mL (Figure 1E), the *NeuroD1* expression decreased in a Dox concentration-dependent manner (Figure 1E), suggesting that a low level of NeuroD1 expression cannot effectively induce MtN conversion but that NeuroD1 *per se* is able to do so if highly expressed. To compare the protein expression levels of NeuroD1 per cell under different Dox concentrations, microglia were transduced with FLAG-tagged NeuroD1 and treated with Dox at 1 μg/mL or 0.01 μg/mL. We observed reduced protein expression of both NeuroD1 and EGFP at single-cell resolution at the lower dose relative to the higher one (Figure 1F). These data indicate that NeuroD1 expression above a certain threshold level is required for the effective induction of MtN conversion.

**Figure 1.**
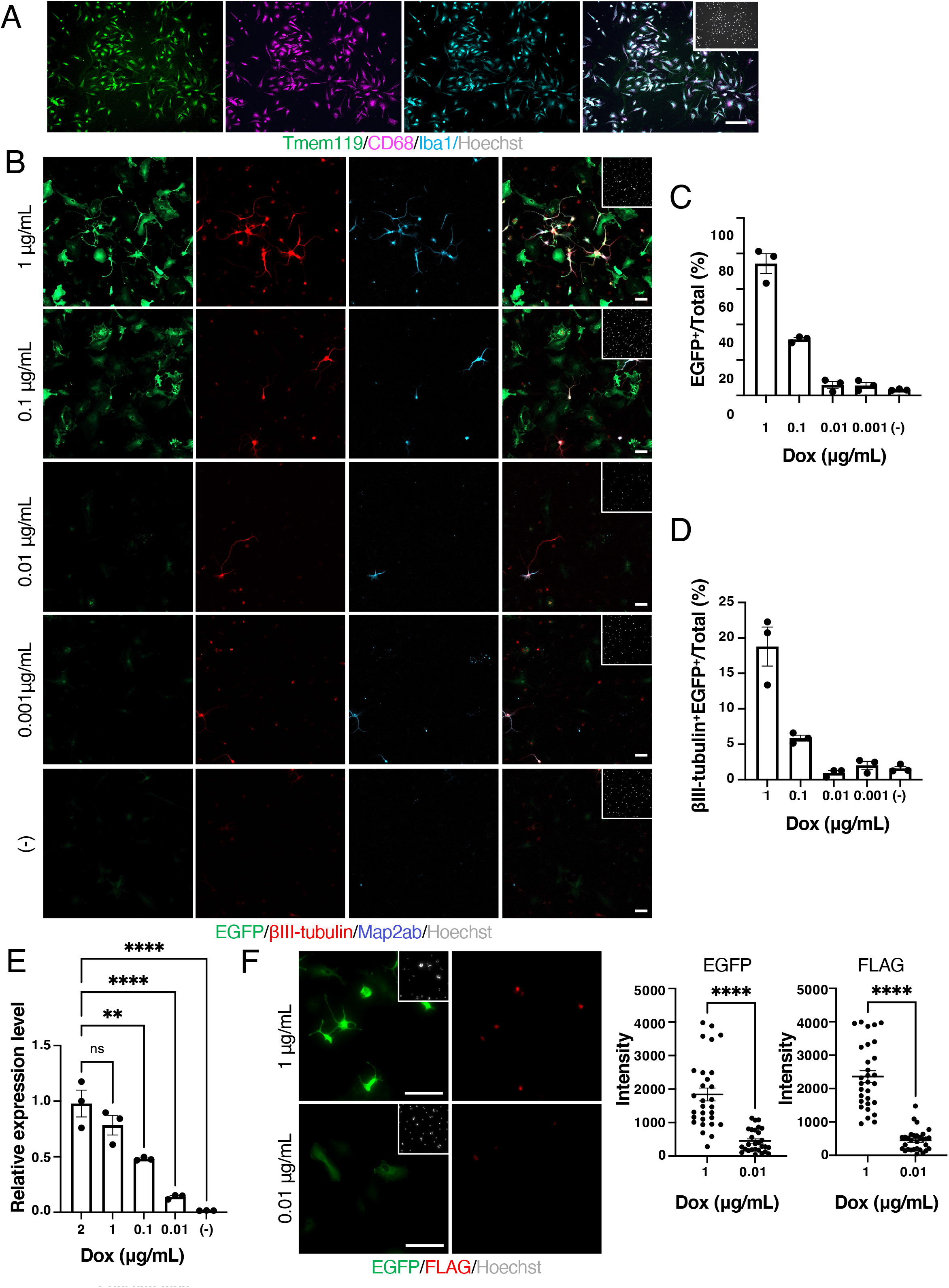
RF expression threshold is required to convert microglia efficiently into neurons. (A) Representative images of staining for microglial markers Tmem119 (green), CD68 (magenta), and Iba1 (cyan) in primary isolated microglia. Scale bar, 75 μm. (B) Representative images of staining for EGFP (green), and the neuronal markers βIII-tubulin (red) and Map2ab (cyan), in NeuroD1-transduced microglia at 7 dpt under the indicated Dox concentration treatment conditions. Scale bars, 50 μm. (C) Quantification of the EGFP^+^ cells in (B) (n = 3). (D) Quantification of the βIII-tubulin^+^ EGFP^+^ cells in (B) (n = 3). (E) qRT-PCR analysis of total *NeuroD1* mRNA levels in NeuroD1-transduced microglia at 2 dpt under the indicated Dox concentration treatment conditions (n = 3 biological replicates). **p < 0.005, ****p < 0.0001 by ANOVA with Tukey *post hoc* tests. (F) Representative images of staining for EGFP (green) and FLAG (red) in FLAG-NeuroD1-transduced microglia at 2 dpt under 1 μg/mL and 0.01 μg/mL Dox induction (left). Intensity of EGFP or FLAG in left panel (right). ****p < 0.0001 by unpaired Student’s t test. ns means not significant (P >0.05).

### Elevated expression level of NeuroD1 enhances neuronal reprogramming

We next explored whether increasing the expression level of NeuroD1 promotes neuronal reprogramming from astrocytes in addition to microglia. We prepared mouse NR-astrocytes *in vitro*, treated them with AraC, and allowed them to grow without growth factors: these astrocytes have a state closely resembling that of astrocytes under the physiological conditions (Figure 2A) (Laywell et al., 2000; White et al., 2011). GFAP-expressing NR-astrocytes were transduced with FLAG-NeuroD1 and EGFP by single (×1) or repeated (×3) viral infections, and we found that the repeated infection increased both NeuroD1 and EGFP protein expression levels at the single-cell level (Figures 2B and 2C). When we checked reprogramming efficiency, single NeuroD1 lentivirus infection hardly induced neuronal reprogramming from NR-astrocytes in agreement with our previous reports (Brulet et al., 2017; Matsuda et al., 2019), whereas elevated expression of NeuroD1 by repeated infections dramatically promoted astrocyte-to-neuron (AtN) conversion at 7 dpt (Figures 2D and 2E). Repeated viral infection (×2) also increased MtN reprogramming efficiency (Figures 2F and 2G). These data indicate that increased NeuroD1 expression enables efficient neuronal reprogramming even from cells that are difficult to convert.

**Figure 2.**
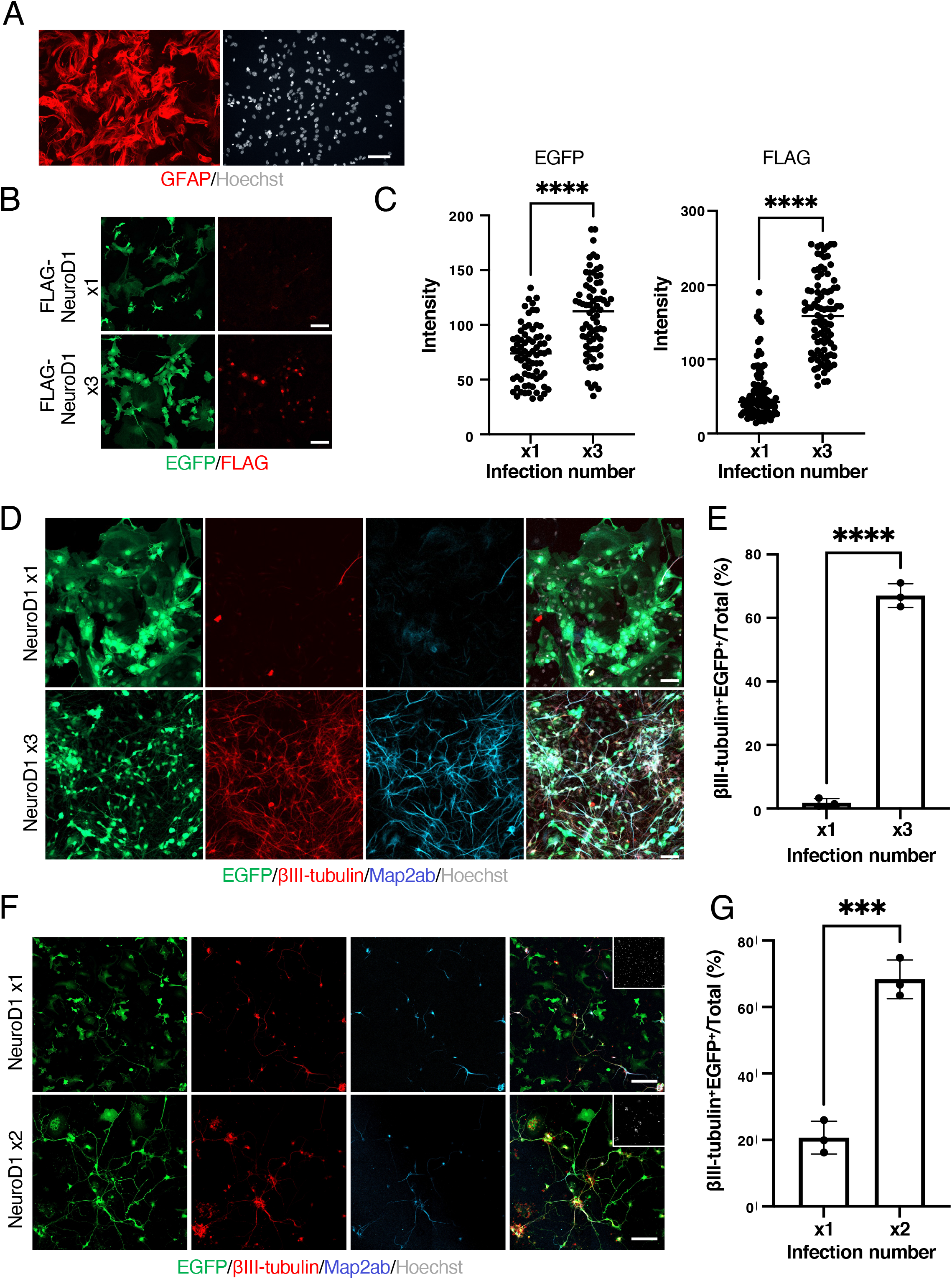
Conversion efficiency is promoted with increased RF expression level. (A) Representative images of non-reactive astrocytes (NR-astrocytes) stained for the astrocyte marker GFAP (red). Scale bar, 75 μm. (B) Representative images of staining for EGFP (green) and FLAG (red) in FLAG-NeuroD1-transduced NR-astrocytes at 2 dpt. Dox, 1 μg/mL. Scale bar, 100 μm. (C) Fluorescence intensity of EGFP or FLAG in (B). ****p < 0.0001 by unpaired Student’s t test. (D) Representative images of staining for EGFP (green), βIII-tubulin (red), and Map2ab (cyan) in reprogrammed neuronal cells from NR-astrocytes at 7dpt. Dox, 1 μg/mL of Dox. Scale bars, 100 μm. (E) Quantification of the βIII-tubulin and EGFP^+^ cells in (D) (n = 3). ****p < 0.0001 by unpaired Student’s t test. (F) Representative images of staining for EGFP (green), βIII-tubulin (red), and Map2ab (cyan) in reprogrammed neuronal cells from microglia at 7 dpt. Dox, 1 μg/mL. Scale bars, 100 μm. (G) Quantification of the βIII-tubulin and EGFP^+^ cells in (F) (n = 3). ***p < 0.0005 by unpaired Student’s t test.

### Differences in cell context affect neuronal reprogramming efficiency

In contrast to our study, astrocytes were previously cultured in the presence of growth factors, FGF2 and EGF (Guo et al., 2014; Heinrich et al., 2010), both of which are expressed in reactive astrocytes and allow the response to signaling pathways critical to neuronal fate choice (Buffo et al., 2008; Burda and Sofroniew, 2014; Robel et al., 2011). To ask whether fundamental environmental differences affect neuronal reprogramming efficiency, we isolated astrocytes and cultured them in the continuous presence of FGF2 and EGF. By this growth factor treatment alone, a small percentage of βIII-tubulin^+^ cells among control virus-infected cells appeared (Figures 3A and 3C), probably because FGF2 and EGF can confer stem cell-like properties on astrocytes, enabling them to differentiate into neurons in accordance with previous reports (Kleiderman et al., 2016; Magnusson et al., 2020). In addition, unlike in the absence of these growth factors (Figures 2C and 2D), we found that even a single NeuroD1 virus infection together with FGF2 and EGF could effectively induce AtN conversion by 7 dpt (Figures 3B and 3C). We then assessed whether subsequent FGF2 and EGF stimulation affect neuronal reprogramming from NR-astrocytes established in the absence of the growth factors. After 3 days’ stimulation with FGF2 and EGF, NeuroD1 expression was induced with 1 μg/ml of Dox in NR-astrocytes infected only once with NeuroD1 lentivirus. We found that FGF2 and EGF stimulation allowed NR-astrocytes exposed to a single NeuroD1 virus infection to be converted into neurons (Figures 3D and 3E). In addition to FGF2 and EGF, LIF is expressed in reactive astrocytes and is known to affect their properties (Linnerbauer and Rothhammer, 2020). However, LIF stimulation did not improve the neuronal conversion efficiency of NR-astrocytes. Thus, these results indicate that environmental factors, especially those that confer stem cell-like properties on astrocytes, can contribute to efficient AtN conversion.

**Figure 3.**
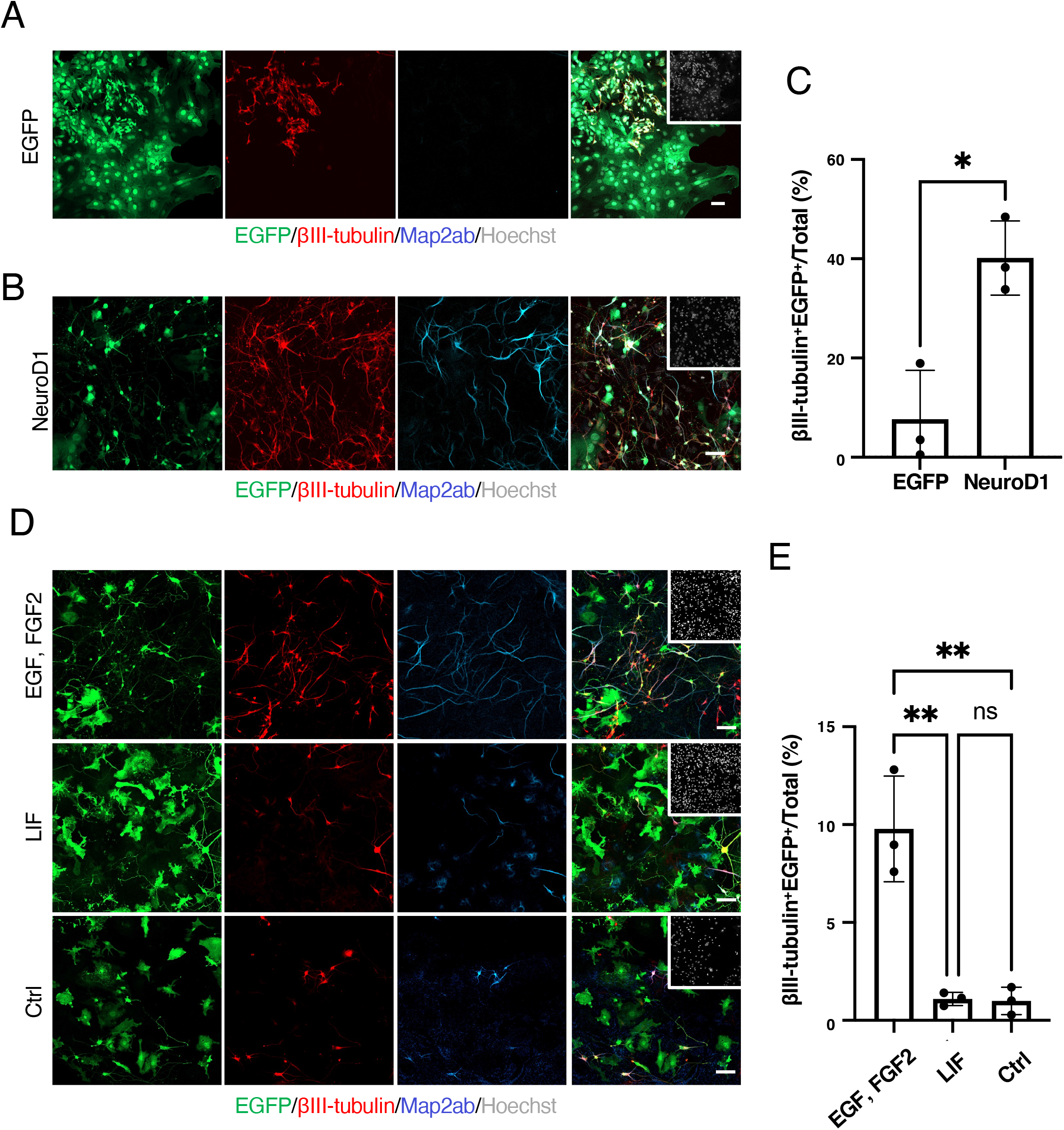
Environmental stimulation affects neuronal reprogramming efficiency. (A) Representative images of staining for EGFP (green), βIII-tubulin (red), and Map2ab (cyan) in EGFP-transduced astrocytes cultured with EGF and FGF2 at 7 dpt. Dox, 1 μg/mL. Scale bar, 100 μm. (B) Representative images of staining for EGFP (green), βIII-tubulin (red), and Map2ab (cyan) in NeuroD1-transduced astrocytes cultured with EGF and FGF2 at 7 dpt. Dox, 1 μg/mL. Scale bar, 100 μm. (C) Quantification of the βIII-tubulin and EGFP^+^ cells in (A and B) (n = 3). *p < 0.05 by unpaired Student’s t test. (D) Representative images of staining for EGFP (green), βIII-tubulin (red), and Map2ab (cyan) in NeuroD1-transduced NR-astrocytes cultured with EGF and FGF2, LIF, and without these factors (Ctrl) at 7 dpt. EGF and FGF2 or LIF were applied for 3 days. Dox, 1 μg/mL. Scale bars, 100 μm. (E) Quantification of the βIII-tubulin and EGFP^+^ cells in (D) (n = 3). **p < 0.005 by ANOVA with Tukey *post hoc* tests. ns means not significant (P > 0.05).

### Combinatorial expression of RFs enhances neuronal reprogramming

Besides increasing the expression level of RFs and environmental factors, combinatorial expression of RFs is another strategy that should be considered to enhance neuronal reprogramming efficiency (Matsuda and Nakashima, 2021). The neurogenic transcription factors Ascl1 and Brn2 have been shown to induce neuronal reprogramming from somatic cells, including astrocytes and microglia (Gascón et al., 2016; Matsuda et al., 2019). Therefore, we first expressed Ascl1 and Brn2 together with NeuroD1 (NAB) in microglia and found that this NAB combination augmented MtN conversion compared to NeuroD1 alone even with a low dose (0.1 μg/mL) of Dox (Figures 4A and 4B). We further observed that the efficiency of AtN conversion was dramatically increased by Dox (1 μg/mL)-induced NAB expression compared to NeuroD1 expression alone, in which AtN conversion was negligible (Figures 4C and 4D). These results indicate that reprogramming efficiency can be positively modulated by combining optimal RFs even under conditions where reprogramming occurs inefficiently.

**Figure 4.**
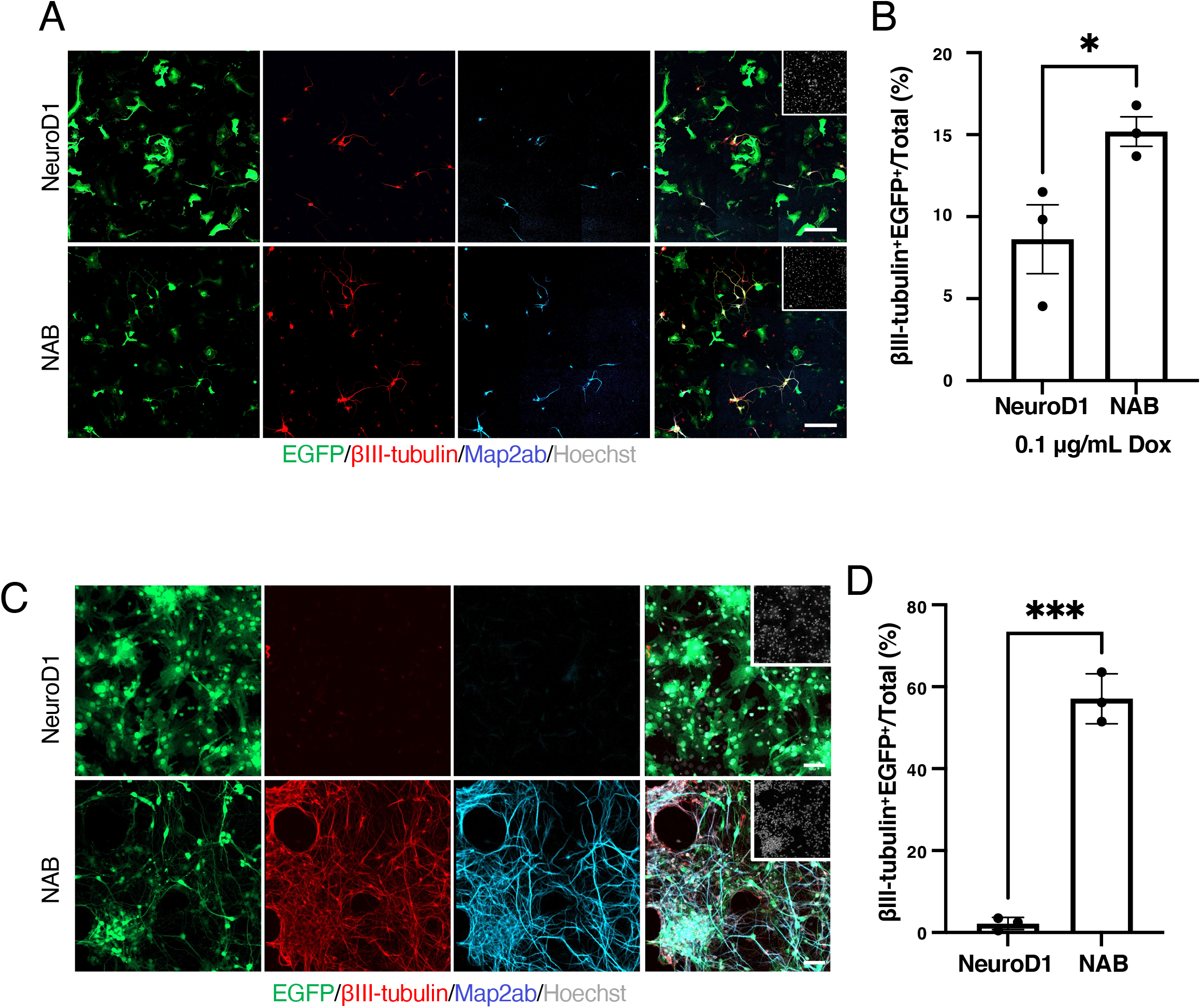
Optimal combinations of RFs increase neuronal reprogramming efficiency. (A) Representative images of staining for EGFP (green), βIII-tubulin (red), and Map2ab (cyan) in NeuroD1- or NAB-transduced microglia at 7 dpt. Dox, 0.1 μg/mL. Scale bars, 100 μm. (B) Quantification of the βIII-tubulin and EGFP^+^ cells in (A) (n = 3). *p < 0.05 by unpaired Student’s t test. (C) Representative images of staining for EGFP (green), βIII-tubulin (red), and Map2ab (cyan) in NeuroD1- or NAB-transduced NR-astrocytes at 7 dpt. Dox, 1 μg/mL. Scale bars, 100 μm. (D) Quantification of the βIII-tubulin and EGFP^+^ cells in (C) (n = 3). ***p < 0.0005 by unpaired Student’s t test.

## Discussion

In this study, using NeuroD1 as a representative RF, we have shown that reprogramming efficiency is influenced by the three factors: RF expression level, environmental factors, and the combination of RFs. In addition, we demonstrated that if the RF expression is sufficiently high, neurons can be induced with a single RF even in cells that are difficult to reprogram. In other words, this finding suggests that the key determinant of successful neuronal reprogramming among the three factors is the RF expression level.

We have previously revealed that in microglia, ectopically expressed NeuroD1 binds to closed chromatin with bivalent modifications, namely active (trimethylation of histone H3 at lysine 4 [H3K4me3]) and repressive (H3K27me3) marks, to induce the expression of neuronal genes (Matsuda et al., 2019). In contrast to microglia, NR-astrocytes lack such bivalent signatures and exhibit a monovalent repressive modification (H3K27me3) around neuronal gene loci, in accordance with the low capacity of NeuroD1 to induce neuronal conversion of these cells (Matsuda et al., 2019). However, another study demonstrated that NeuroD1 could occupy loci possessing the H3K27me3 modification to initiate neuronal programs in ES cells (Pataskar et al., 2016). In the present study, we found that a relatively higher expression level of NeuroD1 induced by repeated virus infections could achieve neuronal reprogramming efficiently from NR-astrocytes. These findings suggest that while NeuroD1 preferentially binds to regions with bivalent modifications, an excess amount of NeuroD1 may increase the likelihood that it will also bind to regions with a monovalent repressive modification to initiate the neuronal program.

Recent studies have shown that the AtN conversion efficiency differs depending on the brain region in which the astrocytes reside. For example, astrocytes in the corpus callosum cannot be reprogrammed into neurons by expression of either NeuroD1 or the combination of Neurog2 and Nurr1, whereas astrocytes in the cortex can be (Liu et al., 2020; Mattugini et al., 2019), implying that the particular environment in different brain regions dictates distinct astrocytic properties and consequently affects reprogramming potential. Astrocytes have been reported to acquire a variety of phenotypes and gene expression patterns in response to many pathological stimuli, such as stroke, neurodegenerative diseases, and aging (Matias et al., 2019). We found in the present study that FGF2- and EGF-stimulated astrocytes are more likely to be converted into neurons than LIF-stimulated astrocytes, although all three of these factors are expressed and regulate the behavior of reactive astrocytes in pathological conditions (Linnerbauer and Rothhammer, 2020). This result indicates that reprogramming efficiency from astrocytes may vary depending on brain pathologies as well as brain regions. Moreover, microglia have also recently been shown to manifest phenotypic heterogeneity across different regions and under neurological diseases in the brain (Deczkowska et al., 2018; Tan et al., 2020). Therefore, it is critical that neuronal reprogramming should be achieved by ensuring sufficient expression of RFs and, if necessary, examining their combinations to apply this technology to brain injury and disease therapy.

Our findings provide insights into how RF expression levels affect neuronal reprogramming efficiency and ways to efficiently induce neurons from two glial cell types, microglia and astrocytes. Boosting reprogramming efficiency should offer therapeutic strategies for neurological conditions such as Alzheimer’s disease, spinal cord injury and ischemia.

## Experimental procedures

### Isolation and culture of primary microglia and astrocytes

We prepared primary microglia and astrocytes from mouse at postnatal day 1 using a previously reported protocol (Matsuda et al., 2019), with some modifications. We dissected cortexes of ICR mice after peeling of meninges to obtain microglia and astrocytes from glial cell mixtures. Dissected tissues were digested with papain (22.5U/ml, Sigma) at 37°C for 30 min and treated with DNase (200U/ml, Sigma). After centrifugation (200 × *g*, 5 min), the cell pellet was suspended in alpha minimum essential medium (MEM) with 5% fetal bovine serum (FBS) and 0.6% glucose and filtered with a 40-μm cell strainer (BD Falcon). After centrifugation (200 × *g*, 5 min), the cell pellet was again suspended in alpha MEM and re-centrifuged (200 × *g*, 5 min). The cell pellets were resuspended in Dulbecco’s modified Eagle’s medium (DMEM)/Ham’s F-12 (Nacalai Tesque) containing 20% FBS, 1 mM L-sodium pyruvate, and MEM nonessential amino acids solution, and treated with GM-CSF (2.5 ng/mL; R&D Systems) to enhance microglial proliferation. This isolated glial mixture was plated in T75 tissue culture flasks (BD Falcon) and the medium was changed every 2–3 days. Subsequently, we collected microglia by strong shaking for 1 h after 7–10 days in culture. The microglia were then plated onto an uncoated 35-mm culture dish and oligodendrocytes were removed by changing the medium 30 min after plating. We used the cells attached to the dish as primary cultured microglia and maintained them in DMEM/Ham’s F-12 containing 20% FBS, 1 mM L-sodium pyruvate, and MEM nonessential amino acids solution.

After isolation of microglia, flasks were treated with AraC (5 μM) for 2 days to remove proliferating cells. Cultures were shaken for 16 h, and then trypsin–EDTA solution was added to the flask to obtain NR-astrocytes. Isolated NR-astrocytes were plated onto an uncoated 35-mm culture dish and maintained with DMEM/Ham’s F-12 containing 20% FBS.

To isolate FGF2- and EGF-stimulated astrocytes, the cell pellet obtained from dissected cortical tissues was cultured in T75 tissue culture flasks using DMEM/Ham’s F-12 containing 20% FBS, hEGF (10 ng/mL, Peprotech), hFGF2 (10 ng/mL, Peprotech), and B27. The medium was changed every 2 days. After 5–7 days of culture, trypsin– EDTA solution was added to the flask and the isolated cells were plated onto an uncoated 35-mm culture dish. FGF2 and EGF (10 ng/mL) or 50 ng/mL LIF were used to stimulate NR-astrocytes.

### Virus production

Lentiviruses were produced by transfecting HEK293T cells in a 10-cm dish with the constructs pCMV-VSV-G-RSV-Rev and pCAG-HIVgp using polyethylenimine. Since lot-to-lot variation in FBS preparations added to the culture medium critically influences the resultant viral tropism, we avoided using FBS for virus preparation (Torashima et al., 2007). After transfection, we cultured the cells with 5 mL of serum-free N2 medium (DMEM/F12 supplemented with insulin (25 μg/mL), apo-transferrin (100 μg/mL), progesterone (20 nM), putrescine (60 μM), and sodium selenite (30 nM)) for 2 days. The supernatant was collected and used for virus infection experiments after filtration through a 0.2 filter to remove cell debris.

### Induction of neuronal conversion

To induce neurons from glial cells, we used lentiviral vectors (derived from the Tet-O-FUW vector) in which gene expression is controlled by the tetracycline operator. Plasmids used in this study are similar to those described in our previous report (Matsuda et al., 2019). For cells to be infected efficiently with the lentivirus, the virus must be added as soon as possible after plating the microglia. Therefore, virus suspensions were added at the time of medium exchange 30 min after isolation of primary microglia, and infection was performed overnight. The medium was then replaced with a neuronal medium (Neurobasal Medium (GIBCO) supplemented with B27 (Gibco), GlutaMAX (2 mM, Gibco), BDNF, GDNF, NT3 (10 ng/mL each, Peprotech), and penicillin/streptomycin/fungizone (Hyclone), and Dox induction was started for 7 days to convert microglia into neuronal cells. Dox was added only once to the medium to activate RF expression. The medium was changed every 2–3 days for the duration of the culture period.

For conversion into neuronal cells from astrocytes, the virus suspension was added at the time the cells were seeded, and infection was performed overnight. The medium was replaced with neuronal medium the next day, and Dox induction was started for 7 days to convert astrocytes into neuronal cells.

For sequential viral infections, a second infection was performed 8 h after the first infection was completed, and overnight virus infection and medium replacement were repeated. To convert glial cells into neuronal cells, the medium was replaced with neuronal medium containing Dox after the final infection and the cells were cultured for 7 days. The medium was changed every 2–3 days for the duration of the culture period.

### Immunochemistry

Cells were fixed in 4% paraformaldehyde for 10 min and blocked for 1 h at room temperature (RT) with blocking buffer (5% FBS and 0.3% Triton X-100). After blocking, the cells were incubated with the following primary antibodies for 2 h at RT: anti-βIII-tubulin (1:500, Covance), anti-Map2ab (1:500, Sigma), anti-GFP (1:500, Aves), anti-FLAG (1:500, Sigma), anti-Tmem119 (1:500, Abcam), anti-CD68 (1:500, Bio-Rad), and anti-Iba1 (1:500, Abcam). Stained cells were visualized with a fluorescence microscope (Axiovert 200M, Zeiss) and a confocal microscope (LSM800, Zeiss). Fluorescence intensity of the cell soma was quantified using LAS AF (Leica) or ZEN (Zeiss).

### Real-time qRT-PCR

Total RNA was isolated using an RNeasy Micro Kit (QIAGEN) according to the supplier’s protocol. RNA quality was checked with a spectrophotometer. Reverse transcription reactions were performed using a SuperScript VILO cDNA Synthesis Kit (Life Technologies) following the supplier’s protocol. qRT-PCRwas performed with SYBR green fluorescent dye using Step One Plus (Applied Biosystems) and Mx3000 (Stratagene). GAPDH was used as an endogenous control to normalize samples. The PCR primers used in this study were NeuroD1_Fw:AAGCCACGGATCAATCTTCTC and NeuroD1_Rv:CGTGAAAGATGGCATTAAGCTG.

### Statical analysis

Data were analyzed using Prism 9 ver.9.1.2. Unpaired Student’s t tests were used to calculate the p value for pairwise comparisons. For multiple comparisons, p values were calculated using one-way ANOVA with the Tukey *post hoc* test. Data represent mean ± SEM. We considered probabilities of p < 0.05 to be significant.

## Author contributions

K.M-I., T.M., and K.N. designed research and analyzed data; K.M-I. and T.M. performed research; K.M-I., T.M., and K.N. wrote the paper.

## Acknowledgments

We thank Y. Nakagawa for excellent secretarial assistance and I. Smith for proofreading the manuscript. This work was supported by a Grant-in-Aid for Research Activity Start-up JP21K20639 (to K.M-I.), a Grant-in-Aid for Scientific Research (B) JP21H02808 (to T.M.), the Takeda Science Foundation (to T.M.), a research grant from the Noguchi Institute (to T.M.),AMED JP21bm0404057 (to T.M., K.N.), and a Grant-in-Aid for Scientific Research on Innovative Areas 17 JP16K21734 (to K.N.).

